# Newly developed *ad hoc* molecular assays show how eDNA can witness and anticipate the monk seal recolonization of central Mediterranean

**DOI:** 10.1101/2021.02.13.431078

**Authors:** Elena Valsecchi, Emanuele Coppola, Rosa Pires, Andrea Parmegiani, Maurizio Casiraghi, Paolo Galli, Antonia Bruno

## Abstract

The monk seal *Monachus monachus* is the most endangered pinniped worldwide and the only one found in the Mediterranean, where its distribution and abundance have suffered a drastic decline in the last few decades. Data on its status are scattered due to both its rarity and evasiveness, and records are biased towards occasional, mostly coastal, encounters. Nowadays molecular techniques allow us to detect and quantify minute amounts of DNA traces released in the environment (eDNA) by any organism. We present three qPCR-assays targeting the monk seal mitogenome. The assays were soundly tested on an extensive and diversified sample set (n=73), including positive controls from Madeira breeding population collected during the peak of abundance, and two opportunistic Mediterranean eDNA-sample collections (offshore/coastal) from on-going projects. Monk seal DNA was detected in 47.2% and 66.7% of the samples collected in the Tyrrhenian from a ferry platform (2018-2019) and in the Pelagie archipelago -Strait of Sicily- (2020) respectively, anticipating (up to 2 year) visual observations occurred subsequently in proximity of the sampled areas. In the Tyrrhenian, detection occurrence increased between 2018 and 2019. Monk seal DNA recoveries were commoner in night-time ferry-samples, suggesting nocturnal predatory activity in pelagic waters. The proposed technique provides a non-invasive and yet highly-sensitive tool for defining the monk seal actual distribution and home range, its recovery rate and pinpoint coastal/offshore localities where prioritizing conservation, research, citizen science and education initiatives.

## INTRODUCTION

The Mediterranean monk seal (*Monachus monachus*) is the only pinniped historically and permanently present in the Mediterranean basin. Originally Mediterranean monk seals were found regularly throughout the Mediterranean, Marmara and Black Seas, along the West African coast to as far south as Cap Blanc, and in Cape Verde, Canary, Madeira and Azores Islands. In the 20^th^ century, the species was drastically reduced, primarily by fishermen, over most of its range (virtually disappearing from Egypt, Israel, Cyprus, Lebanon, Montenegro, the Black Sea, Italy, Corsica, France, the Balearic Islands, Spain, Tunisia, and Morocco), leading to severe bottleneck and inbreeding depression signatures in the refugial populations in eastern Mediterranean (Stoffel *et al*., 2018; Karamanlidis *et al*., 2021). However, in the last decade, occasional sightings on Italian coastal waters have been recorded (Supplementary Fig. S1).

The Mediterranean monk seal current distribution is only partially known with most available data retrieved from sites where adult females are known to return every year to reproduce. Thanks to the predictability of these events, some of the delivery caves are monitored year-round using camera-trap (e.g. Martinez-Jauregui *et al*., 2012), allowing us to learn much about mother/pup relation. Yet, the home ranges of these marine mammals are wide, as adult individuals are known to be able to cover several tens of kilometers per day (Adamantopoulou *et al*., 2011), and little is known about milling and foraging areas and which is the actual distributional range during the non-breeding season. This kind of information is difficult to retrieve as monk seals are often elusive. The recently become available possibility to molecularly monitor the presence of any target species through the analysis of traces of its DNA released in the environment (eDNA) (Beng and Corlett, 2020; Taberlet al., 2012) provides a precious and absolutely non-invasive means by which unveiling and oversee the Mediterranean monk seal population.

Here, we make available a triple set of species-specific molecular assays able to detect the presence of residual Mediterranean monk seal DNA, both from soil and water environmental matrices. The developed assays were validated both in silico and in the wet lab on a wide range of DNA templates, including tissue (n=3) and residual (n=12) extracts and marine eDNA samples (n=58). The overall sample (n=73) consist of 7 categories: 3 positive and 2 negative controls and 2 sets (offshore/coastal) of marine eDNA samples. Our results show that 1) the assays are successful in identifying the smallest amount of monk seal eDNA (up to 5.7*10^−8^ mg/L); 2) the screening on both opportunistic Mediterranean marine eDNA sample sets (from previous and ongoing studies) detected the presence of monk seal in about 50% of the samples of either set, and showed that eDNA analysis can anticipate the detection of the monk seal presence in the study areas before human eye; 3) the proposed approach can potentially unveil still unknown aspects of the biology of this charismatic marine mammal species, by allowing its identification in inaccessible contexts (i.e. offshore waters and during night-time); 4) finally, the possibility to monitor the monk seal presence also in offshore waters allows the acquisition of spatial data useful for the identification of areas that achieve quantitative conservation targets (i.e. amount of the species range to be protected).

## MATERIALS AND METHODS

### Species specific primer sets design

Candidate regions for designing monk seal specific primers were identified within the mtDNA regions targeted by MarVer primers (Valsecchi *et al*., 2020), as the amplicons produced by the three primer sets -MarVer1, MarVer2 and MarVer3 - were all highly polymorphic also between the two sister species of the Mediterranean (*Monachus monachus*) and the Hawaiian (*Neomonachus schauinslandi*) monk seals: 12 (out of 199 bp), 14 (out of 87 bp) and 24 (out of 237 bp) variable sites were found in the fragments amplified by MarVer1, MarVer2 and MarVer3 respectively. Internally to each amplicon, one monk-seal specific primer was designed from the stretch of sequence showing the largest number of diagnostic sites. Each monk-seal specific primer was paired for amplification to the corresponding MarVer universal primer (Valsecchi *et al*., 2020) on the opposite strand. In those instances where the universal primer presented one degenerate base, the mammalian variant was used (Valsecchi *et al*., 2020). For the reasons exposed in the Discussion, another prerequisite we included in the search for adequate priming sites was that of aiming at producing amplicons sufficiently different in size between the three loci. Hereafter we use the terms “MarVer1”, “MarVer2” and “MarVer3” to designate the three loci, rather than the primer sets.

### Sample set

The three assays were tested on seven categories of samples, including both tissue/biological residues DNA (n=3) and marine eDNA (n=4) templates, encompassing three kinds of positive controls, two negative controls and two trials on Mediterranean marine eDNA “opportunistic” samples already available from ongoing projects from EV’s research group (Table 1). The seven categories are as follow:

**Table 1.**
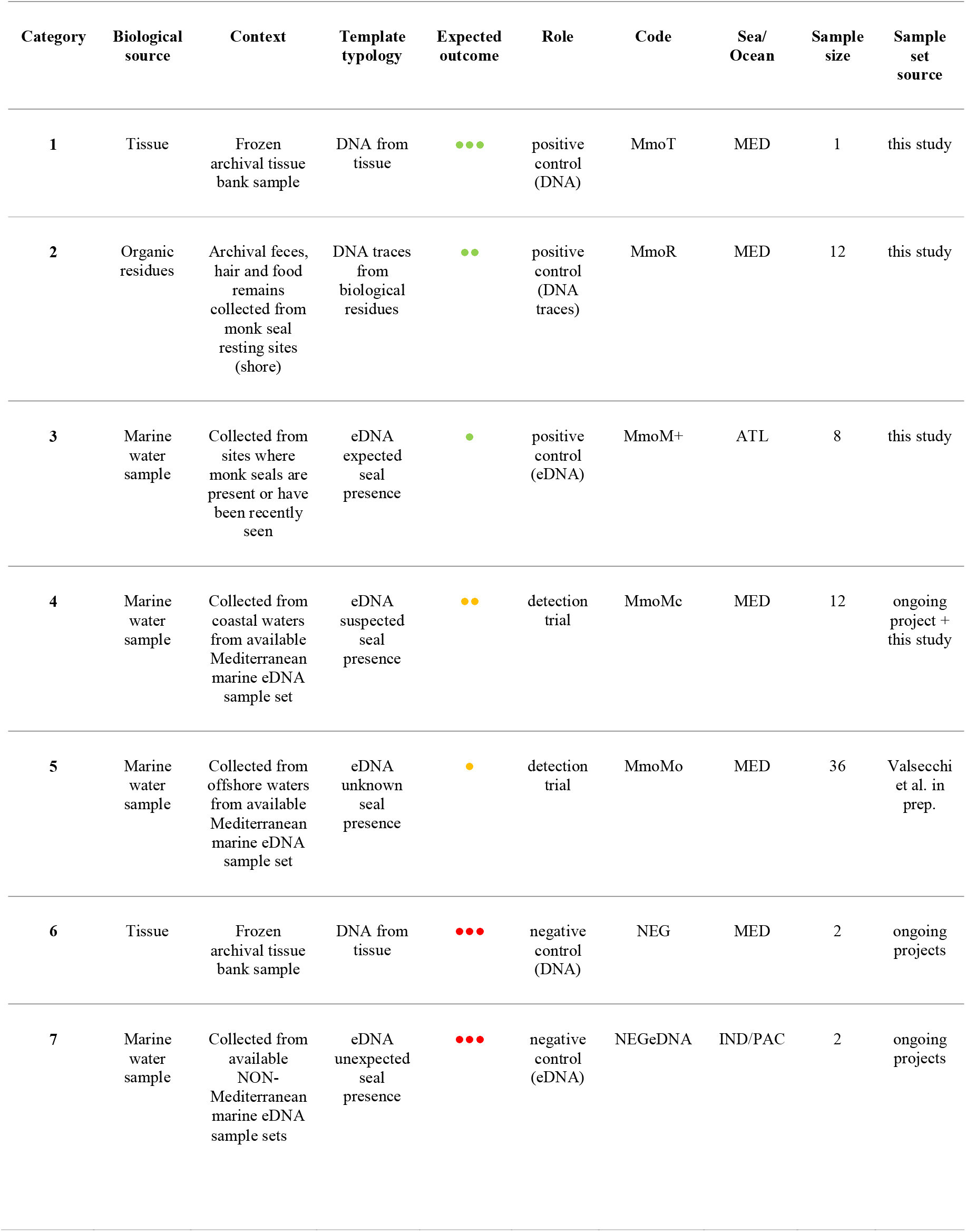
Samples categories. In the “Expected outcome” column, green, yellow and red dots indicate sample categories where the monk seal presence is expected (positive controls), unknown but possible (Mediterranean trial samples) and impossible (negative controls), respectively. The number of dots reflects strength of expected outcome

#### Category 1 <u>DNA positive control</u>

DNA extracted from tissue samples (spleen, sample #474, Marine Mammal Tissue Bank of the University of Padua, www.marinemammals.eu);

#### Category 2 <u>DNA-traces positive control</u>

DNA extracts from 12 environmental residual samples (hair, feces, food remains and regurgitations) referable to 8 circumstances (1 to 3 samples per instances) in which monk seals remnants were collected by EC in known monk seal sites (7 in Favignana and 5 in Croatia) between 2011 and 2014, and stored in ethanol or as dried samples (Supplementary Table S1);

#### Category 3 <u>eDNA positive control</u>

The most suitable area from where obtaining marine eDNA samples likely to contain monk seal eDNA, was identified in the Madeira archipelago, where an isolated population of monk seals (estimated 20-30 individuals) is found and has been protected and monitored for over 30 years (Pires et al. 2008). We waited for the month of November when the breeding season is at its apex and the number of newborn pups is at its maximum (Pires et al. 2008), thus ensuring high seals’ activity in proximity to coastal sheltered shores and caves. Eight marine eDNA samples were extracted from marine water samples (each of 8-12 L) collected in Madeira (Desertas Northern Island n=7 and Madeira main island n=1) during the 2020 breeding season (10^th^ −12^th^ of November, Supplementary Table S1). The eight samples were purposely collected in points with variable incidence of seal presence: ranging from areas with no recorded seal to points in proximity of the entrance of caves currently inhabited by seals (Figure 3). Water sample collection was operated by the rangers of the National Park Service of Madeira and the expected likelihood of finding monk seal molecular signals was initially not shared in order to ensure an unbiased molecular detection. These positive control samples were used not only to assess the detection capability of the developed assays (quality check), but also to calibrate the intensity of the molecular signal (quantity check) in order to allow a more accurate estimate in surveyed surveillance eDNA samples.

We tested the molecular assays also on “real” Mediterranean marine eDNA samples, available from previous and ongoing projects, collected in areas where the presence of the monk seal is considered extremely rare but not impossible, as suggested by a few recent sightings (Supplementary Figure S1), some of which occurred in proximity of our sampling sites, month to years after our sample collection. Thus, the following two groups of eDNA samples were tested:

#### Category 4 <u>Trial on Mediterranean coastal eDNA samples</u>

This subset of samples was made of two components: the first consists of available eDNA samples from one ongoing project (n=10) collected in the water surrounding the Pelagie Islands between the 14^th^ of August and the 25^th^ of September 2020, and two additional samples collected from shore in the days following reported monk seal sightings (22^nd^, 31^st^ of October and 9^th^ of November 2020, Supplementary Figure S1) in proximity of the sighting point in the waters of Lampedusa island, in order to relate the intensity of the signal to the actual presence of seals. Samples details are found in Supplementary Table S1.

#### Category 5 <u>Trial on Mediterranean offshore eDNA samples</u>

These samples were collected in the Northern Tyrrhenian Sea as part of an ongoing project relying on the collection of marine water samples from operating ferries (Valsecchi et al. submitted, “MeD for Med” project). The surveyed route was the Livorno-Golfo Aranci, run by Corsica Sardinia Ferries, during summers 2018 (n=16) and 2019 (n=20). Unfortunately, no 2020 sample was available due to the restrictions imposed by the COVID epidemic. The sampling design involves the collection of water samples from fixed sampling stations (constant over the different cruises) and additional samples gathered on occasion of sightings of cetaceans operated by members of the FLT network team of ISPRA (Arcangeli et al. 2017).

A double set of negative controls was included in our test:

#### Category 6 <u>Tissue-extracted DNA negative controls</u>

Bottlenose dolphin (*Tursiops truncatus*) and yellowfin tuna (*Thunnus albacares*) DNA samples were used as negative controls when processing categories 1 and 2 samples.

#### Category 7 <u>Non-Mediterranean eDNA negative controls</u>

Environmental DNA samples from international ongoing projects (Maldives and Hawaii) were used as *M. monachus* negative controls in environmental samples (categories 3, 4 and 5).

The DNA from the tissue sample (category 1) was extracted using the Qiagen DNeasy Blood and Tissue Kit, while the DNA of biological residues and environmental samples (categories 2-6) was extracted using the Qiagen DNeasy PowerSoil Kit according to the manufacturer protocol. When biological residual samples (MmoR, category 2) were in liquid form (e.g. feces dissolved in ethanol), an aliquot was filtered through a membrane that was subsequently rinsed with distilled water prior DNA extraction. Supplementary Table S1 shows a detailed description of all 73 samples.

The four sets of eDNA samples (categories 3 to 5 and 7), coming from different projects and having been collected in different contexts and by different operators, were not always homogeneous in terms of number of liters of marine water being processed and porosity of filters used. However, they were all collected from the sea surface, with the exception of ferry samples collected from sea water intake placed 4.5 m below sea level. Water volumes were collected and stored in the Bag-in-Box Sampling System (BiBSS) (Valsecchi et al., in prep.).

The two samples collected directly from shore (MmoMc11 and MnoMc12, category 4) underwent through a ten-fold template dilution prior amplification, a procedure used to minimize the interference of PCR inhibitors (Gasparini *et al*., 2020), known to be particular abundant in proximity of sediments (Lance and Guan, 2019).

### qPCR experiment design

Quantitative Real Time PCR (qPCR) assays were performed with AB 7500 (Applied Biosystem) to test the three primer pairs. First, for each primer pair, *M. monachus* tissue samples were used to set amplification efficiency (E), limit of detection (LOD) and limit of quantification (LOQ), according to Bustin *et al*., 2009 and Klymus *et al*., 2019. Ten-fold serial dilutions were used to generate the standard curve.

All 73 samples were run in triplicate, using the following qPCR conditions: an initial denaturation at 95 °C for 10 min, followed by 40 cycles of denaturation at 95 °C for 15 s and annealing-elongation at 56 °C for 1 min. A final dissociation stage was performed. Amplification reaction consisted of 5.0 μl SsoFast EvaGreen Supermix with Low ROX (Bio-Rad), 0.1 μl each [10 μM] primer solution, 2 μl DNA sample, and 2.8 μl of Milli-Q water. For each run, positive controls, tissue/eDNA negative controls, plus no-template negative controls were included in triplicate, as well.

Ct (Threshold Cycles) values were converted into counts (DNA copies) (Bruno *et al*., 2017). When qPCR copy number outputs were below the limit of quantification (LOQ), but above the theoretical limit of qPCR (three copies per reaction according to Bustin *et al*. (2009)), they were addressed as ‘detectable but not quantifiable’ (DBNQ). For all the reactions, primer specificity was verified by the calculated TM.

### Testing for correlation between marker’s amplicon size and DNA degradation

Since the primer set producing the larger amplicon (MarVer3, 216 bp) was found to perform less efficiently than the other two (see Results), we tested whether the drop of signal between a high-quality DNA sample and a degraded DNA sample (as eDNA) was the same for all three loci (whose sizes are multiple of ca 70 bp), regardless the dimension of the fragment they amplify. This was possible thanks to the availability of two exceptional samples in our data set: 1) a high molecular weight DNA sample, such as MmoT01, freshly extracted from a tissue collected in the immediate *post-mortem* and 2) the best possible eDNA sample, such as MmoM+01, collected in a site at high monk seal eDNA concentration. This sample was collected in a site characterized by two peculiar conditions: a) proximity to a cave contextually inhabited by many seals and b) limited mixing with surrounding waters (thus little signal dispersal) due to a counter-current flux pushing the water against the opening of the cave (see Discussion). All three loci produced a strong signal in both samples and all replicas thus allowing to test for correlation between amplicon size and DNA quality. We applied a Kruskal-Wallis test, as not requiring data being normally distributed and having similar levels of variance.

## RESULTS

### Primer sets

Within each of the three MarVer loci (Valsecchi *et al*., 2020) one *Monachus monachus* specific priming site, fulfilling the desirable criteria of high specificity for the target species, was identified (Table 2). The degree of differentiation of the three species-specific priming sites between the Mediterranean monk seal (GenBankAN: GU174602) and its closest relative, the Hawaiian monk seal (GenBankAN: MscNC_008421), was as follows: 6 bp (24%), 9 bp (32.1%) and 8 bp (22.2%) for MarVer1, MarVer2 and MarVer3, respectively. Primers sequences have been deposited on the BOLD System repository (Ratnasingham and Hebert, 2007).

**Table 2.**
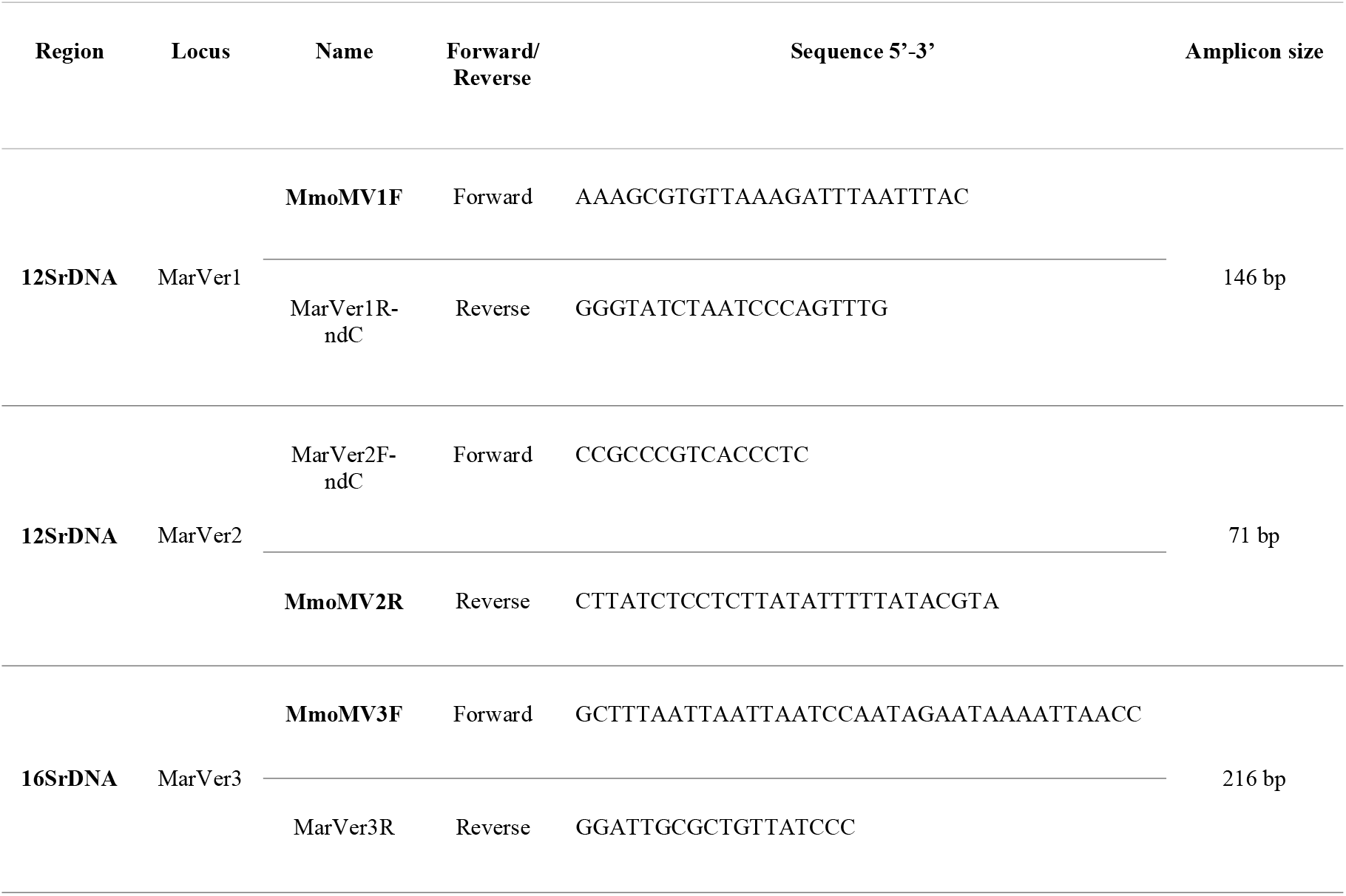
Primer sequences. Primer sets sequences for the three molecular assays specific for *M. monachus*. Oligos named “Mmo…” are the species-specific primers (in bold), while those labelled “MarVer…” are the universal marine vertebrate primers described in Valsecchi et al. (2020). The suffixes “nd” followed by a letter (e.g. “ndC”) indicate the non-degenerate form of a specific primer where the letter indicates the mammalian specific nucleotide used in place of the degenerate base (see Valsecchi et al., 2020).

### qPCR outcomes

A total of 1710 qPCR reactions were run over the 73 samples covering the seven sample categories (Table 1). MarVer1 amplification efficiency reached 92.6%, LOQ corresponded to Ct=34.75; MarVer2 amplification efficiency reached 100.7%, LOQ corresponded to Ct=35.8; MarVer3 amplification efficiency reached 92.2%, LOQ corresponded to Ct=34; R2≥0.99 for each assay. Melting temperatures were 75.4 +/-0.3, 71.5 +/-0.3, 78.5 +/-0.2 °C, for MarVer1, MarVer2, and MarVer3, respectively. According to the LOQ calculated for each locus, qPCR DNA detection outcomes were divided in three classes: 1) no signal, 2) monk-seal DNA detectable but not quantifiable (DBNQ), 3) positive quantifiable detection (PQD). Over the 73 samples, 26 were PQD positives for at least one of the three markers, of which 19 (90.5%) and 7 (14.6%) fell into the positive control group (n=21) and the trial eDNA samples (n=48), respectively (Fig. 1 and Supplementary Table S1). We found a total of 19 DBNQ positives (for at least one of the three loci), of which only 1 (4.7%) in the positive control group, and the remaining 18 (37.5%) in the two Mediterranean trial eDNA samples (2 in the Strait of Sicily and 16 along the ferry track in the Northern Tyrrhenian, see below). Twenty-eight (28) samples gave no signal of monk seal DNA: they included one positive control (eDNA), sample MmoM+02, and all four negative controls (tissue-extracted DNA and eDNA). The remaining 23 samples were Mediterranean eDNA samples, namely 19 (52.7%) and 4 (33.3%) in the Tyrrhenian and in the Strait of Sicily samples respectively (see below).

**Fig.1.**
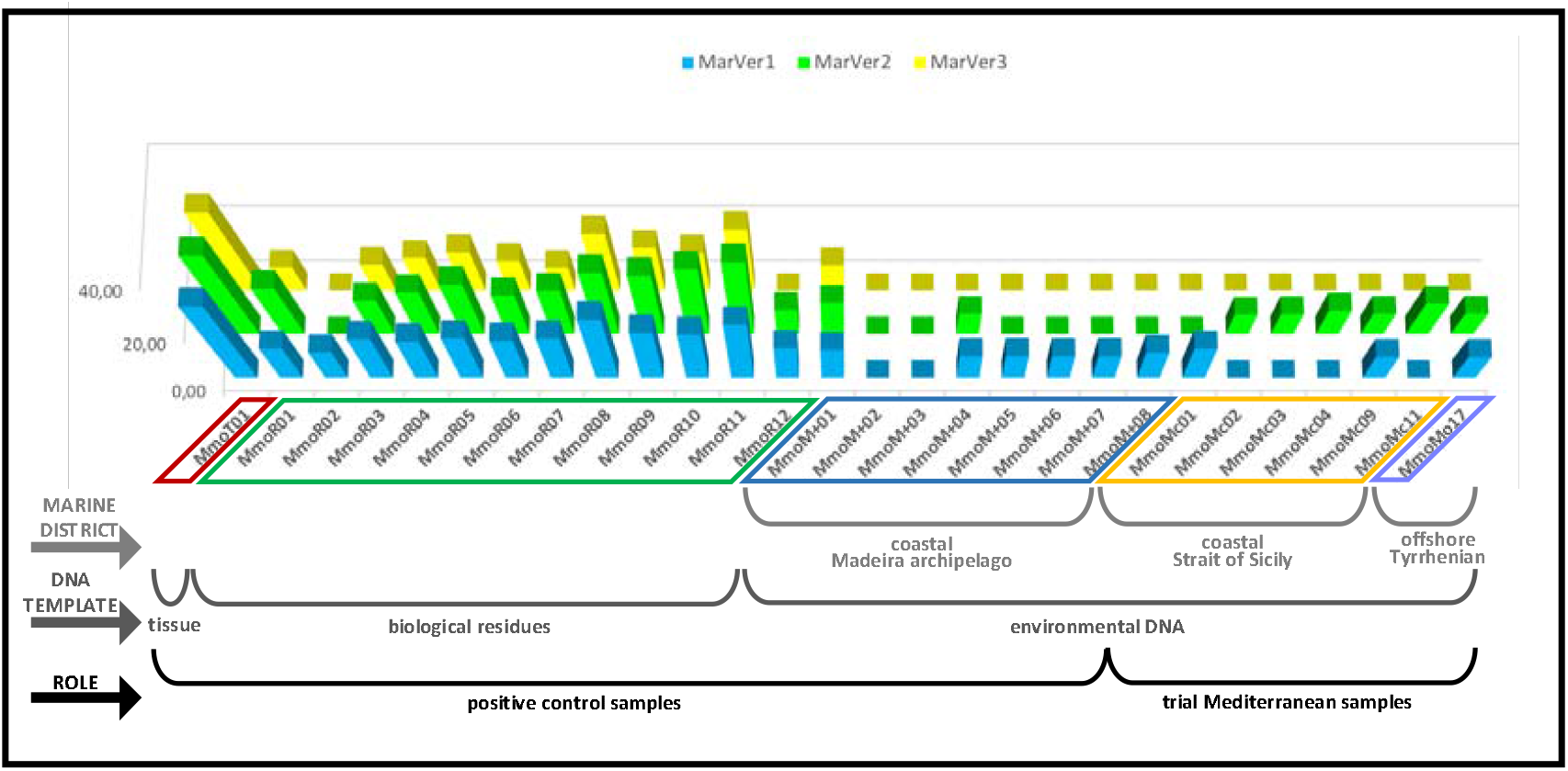
Distribution of Log_2_ DNA-copies in positive control and PQD samples. Log_2_ DNA-copies distribution across positive quantifiable and detectable (PQD) samples. Values (expressed per liter of marine water in the case of eDNA samples) for all three markers across the entire set of positive control samples are shown in comparison with the same values recorded in the 7 Mediterranean eDNA samples (last ones on the right) reporting unambiguous signals of the monk seal presence (i.e. above the detectable and quantifiable threshold).

Samples being positive for monk seal DNA for all three loci (and sample/experimental replicas) were only found within the positive control group (Categories 1, 2 and 3). Most environmental DNA samples (including 5 of the 8 positive control eDNA samples from Madeira) show positiveness to monk seal DNA above the LOD with only one or two of the three loci (Fig. 1, Supplementary Table S1). The locus recovering monk seal DNA in more samples (n=22) was MarVer1 locus, followed by MarVer2 (n=20). The 16SrDNA primer set, MarVer3, was the one recovering the lower number of positive quantifiable detection (n=12), all of which found in positive-control samples, and only one of these regarding an eDNA sample (MmoM+01). The complete list of Log_2_ DNA copies recovered in the complete data set, for all three sample and experimental replicas and markers, is shown in Supplementary Fig. S2.

### Positive-control samples (categories 1 to 3). Category 1

Monk seal tissue-extracted DNA (MmoT1), with a concentration of 284 ng/μl (NanoDrop 2000, ThermoFisher Scientific), was used as refence sample. For this sample all three primer sets produced amplicons of the expected size in all 3 reaction replicas, allowing the detection of up to 5.7*10^−8^ mg/L. **Category 2** All 12 DNA extracts from biological residues (MmoR samples in Tables 1 and Table S1) produced at least one PQD in the triplicate test with each one of the assays. The number of samples scoring PQD in all three replicas were 11, 10 and 8 for MarVer1, MarVer2, MarVer3 respectively. Thus, within category 2 samples, MarVer1 produced overall more PQDs (34 out or 36) than the other two loci (31/36 and 27/36 for MarVer2, MarVer3 respectively), although MarVer2 produced the highest number of DNA-copies. **Category 3** The results obtained on the positive control eDNA samples collected off the Madeira archipelago and the field observations reported by the rangers at the time of water samples collection are shown in Fig. 2. Sample MmoM+01, collected at the entrance of the Tabaqueiro cave (attended by approximately 8 seals), was the control eDNA sample producing the strongest signal, in all three sample replicas, for all the experimental triplicates and for all three markers (Supplementary Table S1, Fig. 2 and Supplementary Fig. S2).

**Fig.2.**
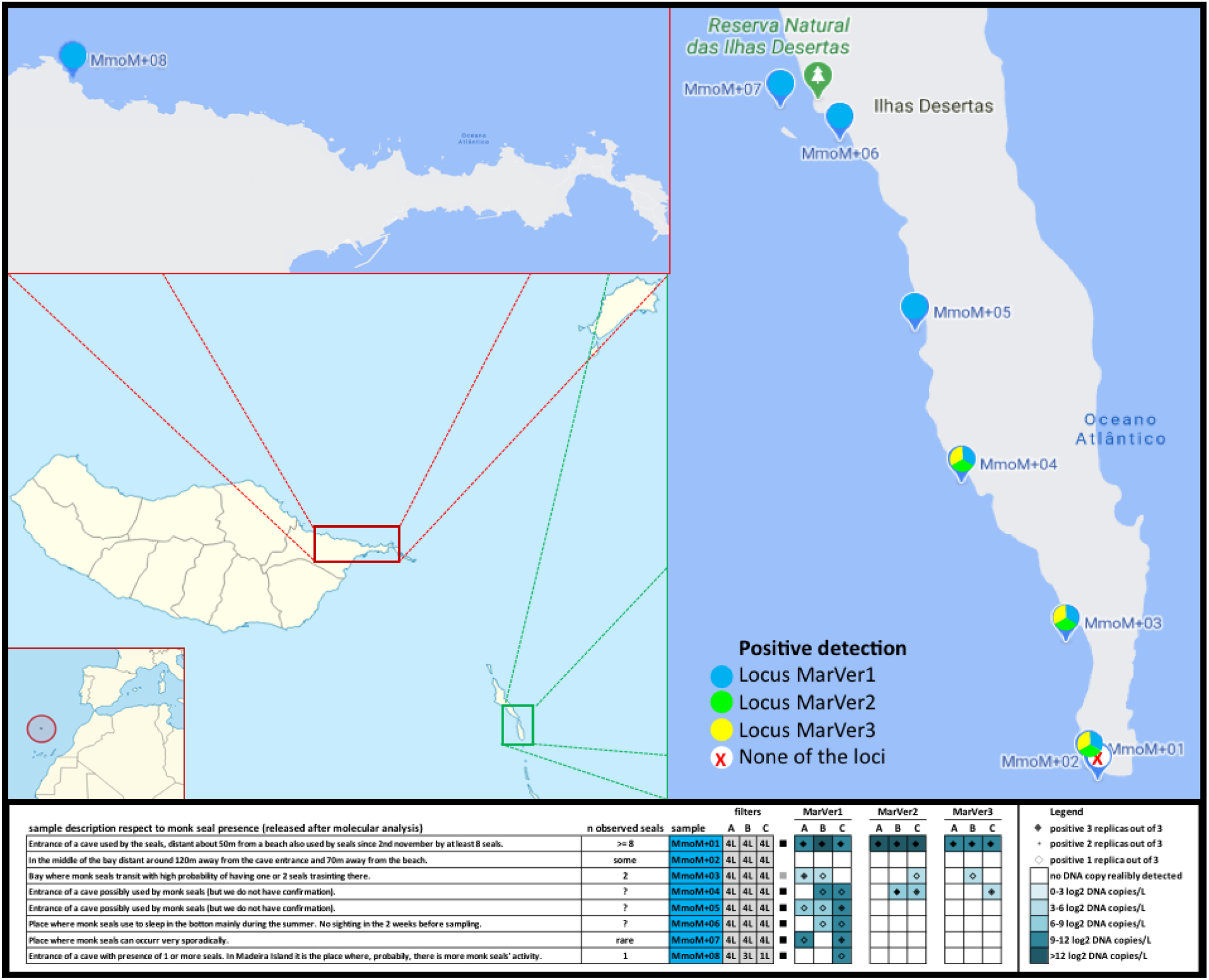
Madeira map, samples and results. Madeira eDNA positive-control (sample category 3) sampling sites and results. In the map the eight sampled sites are indicated with color-coded dots according to which marker detected monk seal DNA in that specific site. The result table reports on the left-hand side the field notes of each sampling site as reported from the rangers at the moment of sample collection and release to the molecular team after disclosure of results. On the right-hand side are shown the molecular results for each of the three assays expressed in ranges of logarithmic scale of number of DNA copies per liter of sampled marine water.

### Mediterranean eDNA sample sets (categories 4 and 5)

Positive quantifiable (PQD, n=7) and unquantifiable (DBNQ, n=18) monk seal DNA detections were found in both Mediterranean eDNA sample sets (Fig. 3). Monk seal eDNA was recovered in 47.2% of the ferry samples (Tyrrhenian, n=36) and 66.7% the coastal samples (Pelagie archipelago, Strait of Sicily, n=12). The proportion between PQR and DBNQ detection was much higher in the Pelagian samples (6 PQD and 2 DBNQ detections) than in the ferry sample set (1 PQD and 16 DBNQ detection).

**Fig.3.**
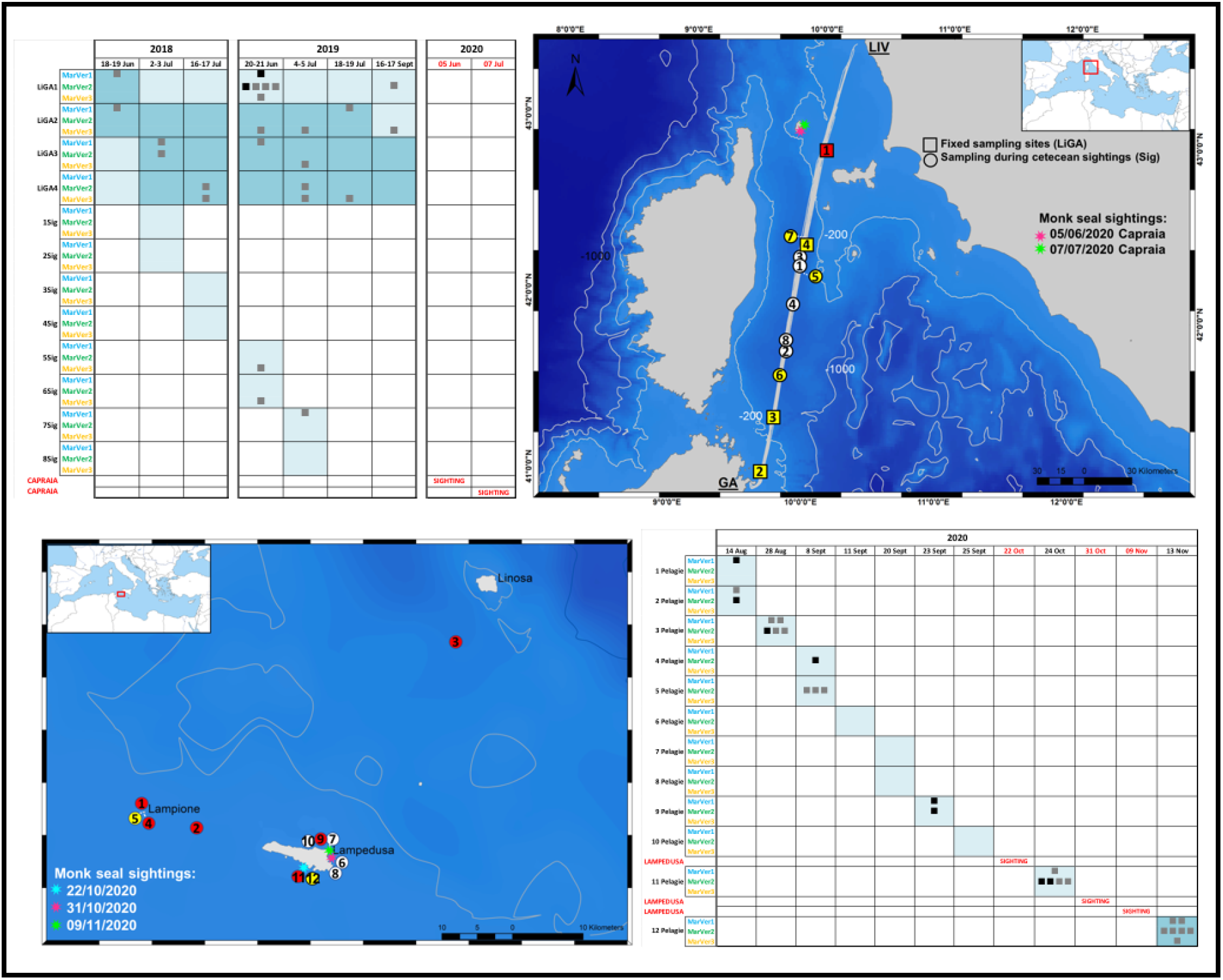
Calendar/map results of Med eDNA samples. Calendar of sample collection and sightings along with molecular results in the two tested Mediterranean areas for which eDNA samples were available from on-going projects: offshore Tyrrhenian (above) and coastal Strait of Sicily, Pelagie archipelago (bottom section). In the tables black squares indicate positive and quantifiable (PQD) monk seal DNA detection by one or more of the three loci (red symbols in the map). The grey squares show samples reporting weak -below the detection threshold-monk-seal signals (yellow symbols in the map), defined in the text as DBNQ (detectable but not quantifiable). The number of squares reflects the number of the sample replicas with positive detection. Coloured cells indicate sampling events, with darker shading showing nocturnal samples. Positions and dates of recent monk seal sightings in the two study areas are reported (in red in the calendar tables).

Excluding the two samples collected in the days following sightings (MmoMc11 and MmoMc12), Pelagian monk seal DNA recoveries (n=6) were more frequent in waters far from the main island of Lampedusa (83.3%, n=5, 4 of which in proximity of Lampione) than in the waters surrounding Lampedusa itself (n=1).

### Negative-control samples (categories 6 and 7)

Both tissue and eDNA negative controls produced no signals in all qPCR runs.

A significant difference was found between the three loci in terms of drop of intensity of the molecular signal (ΔLog_2_ DNA-copies) from the tissue (MmoT01) to the eDNA (MmoM+01) control samples (Kruskal-Wallis test: X^2^= 13.18, df = 2, p-value = 0.001374). The major difference was imputable to locus MarVer3, which showed a significantly higher drop of signal compared with the two 12SrDNA markers, which instead had a similar decrease of signal (Supplementary Fig. S3).

## DISCUSSION

### Assays overview

#### Specificity and sensitivity

The high degree of molecular differentiation of the three presented probes compared to the homologous regions in the sister species -Hawaiian monk seal-genome suggested high specificity of the proposed assays, assumption that was confirmed in all positive and negative control samples sets. High specificity to target species is a fundamental prerequisite for the dissemination of an innovative method given the conservation relevance of the study species. Our experimental design and trial suggest that there is little room for false positives (except those attributable to lab contamination, for which maximum caution should be posed, as in any experiment that is based on eDNA analysis), while incidence of false negative should be more of a concern, as PCR inhibitors present in environmental matrices is recognized as one the main limitations of eDNA-based detection approaches (Loeza-Quintana *et al*., 2020).

#### Reliability

We reckon that also the samples being positive in only 1 or 2 of the three amplification replicas represent genuine monk seal detection, considering the following points:

i. The high homology of the probe to -and only-monk seal DNA (see above), supported by the specific melting temperatures recorded, suggests that any positive detection has to denote monk seal DNA presence.
ii. Failing amplification in replicas of a positive sample may be ascribable to the extremely low concentration of the target molecule. Marine vertebrate DNA counts for only about 0.004% of the eDNA retrieved from marine samples (Stat *et al*., 2017). In the DNA metabarcoding analysis carried out on the ferry sample subset (2018) and same loci, over 90% of the detected vertebrate reads was attributable to bony fish community (decreasing of one more order of magnitude the incidence of rare vertebrates, such as marine mammals, on the overall marine vertebrate fauna), and no read was attributable to monk seal (Valsecchi *et al*., in prep.). Replications (from sampling to DNA amplification) reduces the rate of false negatives; however, the optimal level of replication is difficult to be estimated, since it strongly depends on the detection probability of the target taxon.
iii. With the exception of MmoM+1, collected in proximity of the entrance of the cave most densely populated by monk seals (n=8 ca) and that returned positiveness for all three loci, all remaining Madeira control-eDNA samples (n=7, 87.5%) resulted positive to monk-seal DNA either only in some sample/reaction replicas and/or for only some of the loci (typically the two 12S-rDNA ones), although seals were definitely around in those waters. In some cases, a positive detection was found in only one of the sample replicates (each originating from one of the three filters used to process a single sample). This effect can be more easily explained, considering the specific characteristics of the animal and the eDNA state released (Lacoursière-Roussel and Deiner, 2019; Turner *et al*., 2014). Large animals, such as marine mammals, release in the surrounding water biological traces often consisting of large clusters of cells rather than of single or a few cells (Valsecchi *et al*., 1998). In this way, when the sample is divided into two or more subsamples, it is likely that the cells, being clustered, are not distributed homogeneously among the various subsamples.

Imperfect detection is an unavoidable feature of most data on species presence/absence even during traditional ecological field studies, when individuals and species that are present at one site are not always all detected, and failure in accounting for imperfect detection may result in biased inference. eDNA assays are clearly imperfect, relying on small amounts of degraded DNA, and still laboratory and sequencing costs limit replication (Ficetola *et al*., 2015). However, the high sensitivity and specificity of well-developed eDNA-based approaches can counteract these limitations.

#### Readiness

Quantitative PCR assays allow to obtain quick outcomes as opposed to DNA metabarcoding approach (requiring library preparation and high-throughput DNA sequencing).

### Why three barcode assays

The first motivation for looking at multiple loci was driven by the possibility of picking the best possible candidate, among 3 loci still poorly explored for species detection purposes. Secondarily, we also made sure to draw probes targeting at fragments of different lengths in order to: a) verify whether amplicon size matters when dealing with degenerate templates such as eDNA; b) testing the possibility of multiplexing all or a combination of the 3 barcodes (see below).

Regarding the first point, we do find that the primer set producing the largest amplicon (MarVer3, 216 bp) is in fact the one that amplifies with more difficulty and/or producing fewer DNA copies than the other two assays (MarVer1 and MarVer2, 146 bp and 71 bp, respectively). This is true also when employing high molecular weight DNA as template. The rate of signal drop between a high concentration/quality DNA sample and environmental DNA is significantly higher than that observed in the two markers targeting smaller fragments (Supplementary Fig. S3). If differential drops in recorded DNA signal is correlated to DNA degradation and thus to temporal variables (time since DNA release into the environment) could be explored in future studies.

The fact that the performance of the three markers does not show a consistent ranking across all samples, but that simultaneously it is not randomly distributed, as patterns seem perceptible within eDNA samples categories (e.g. MarVer1 performed better in category-3 samples, MarVer2 in category-4 samples, and MarVer3 was the only marker detecting - weak-signals in some category-5 samples, see Fig. 3, Supplementary Fig. S2, and Supplementary Table S1), suggests further investigations should be encouraged to fill the knowledge gaps about eDNA molecules destiny (Stewart 2019). In Supplementary Fig. S4, we advance a provisional hypothesis of the temporal driven “environmental marker selection”, which attempts to explain the observed phenomena.

Now, our study suggests that the most informative and efficient markers are the two 12SrDNA loci (MarVer1 and MarVer2). MarVer3 (16SrDNA) could be employed in the markers panel and, in case of positive detection, it could indeed -but cautionary-provide a proof of signal freshness (see below in the Mediterranean sample set discussion).

### Traditional PCR and assays multiplexing

The possibility to visualize the assays outcomes using the traditional PCR, rather than qPCR, is a desirable feature, as it allows the test to be carried out in any structure equipped with essential molecular biology laboratory apparatus, besides being cheaper. Although the diagnostic sensitivity decreases in the case of traditional PCR, our study indicates that when the signal is not too weak (see Supplementary Fig. S5), even basic PCR allows the identification of traces of monk seal DNA. The possibility of using the traditional PCR, in turn, entails further advantages, such as the possibility of multiplex run, particularly advantageous when multiple assays are available. This procedure reserves more plusses. Firstly, it allows saving on the amount of eDNA template (precious as typically limited) to be used. Another advantage is the possibility of amplifying the template molecule with different probes at once, a strategy that may reveal useful when dealing with highly diluted eDNA of the desirable target species and especially for large body size species which are likely to shed “clustered” eDNA (e.g. only one of the three replica-filters may contain the target species DNA, although they all come from the same water sample). In these instances, multiplexing could minimize the possibility of false negatives while possibly gathering clues about the signal age (see Supplementary Fig. S4), although the latter aspect needs further deepening. A preliminary multiplex test indicates that all combinations of the three proposed assays are compatible for multiplexing and that the approach is efficient also with eDNA samples, providing a relatively high (say >10 log_2_DNA copies/L) monk seal DNA content such as in Madeira control sample MmoM+01 (Supplementary Fig. S5).

### Currents matter

This study provides indirect evidence of how marine eDNA suffers susceptibility to currents disturbance: Madeira samples stations 1 (MmoM+01) and 2 (MmoM+02) are only 112 m apart from each other, and yet, in the first one (at the entrance of the cave frequented by >8 seals) the molecular signal was the loudest of our eDNA sample set, while in the latter no signal at all was found over 27 reactions (3 filters, 3 replicas, 3 markers). Probably, along those 112 m a current barrier prevents the two water masses from merging (Supplementary Fig. S6). This also implicitly means that around station 1 there is a sort of pocket or relatively unstirred water, where dated and recent monk seal biological traces may accumulate overtime. This fortuitous circumstance makes station 1 the ideal scenario where testing the differential ability of three markers to perceive the temporal scale of eDNA traces. The discrepancy in monk seal eDNA content between the two closely adjacent sampling stations if, on one hand, highlights relevance of currents in swiping away eDNA signals, on the other hand, it adds a reassuring element on the fact that when a signal is indeed intercepted, this probably reflects the presence of the animal in a limited spatio-temporal range (as signals that disperse with currents become shortly extremely diluted and therefore remain undetected).

### Detection of monk seal in Mediterranean eDNA data sets and anticipation on visual data

Positive detections were found in both the Tyrrhenian and the Strait of Sicily trial sample sets, both collected in marine districts where the monk seal is occasionally encountered. DNA recovery was more prevalent in the Pelagian (especially around Lampione Island) than in the ferry sample. Monk seal sightings in waters surrounding the Italian peninsula have been occurring rarely, but constantly in the last decades (Supplementary Fig. S1). According to available published data (some sightings may remain unrevealed for the species safeguard), no sighting was recorded in 2019, while at least nine events were recorded in 2020, suggesting an increased monk seal occurrence in Italian waters. The same trend, but anticipated by a year, was found in our two-year spanning Tyrrhenian sample: the 17 ferry eDNA samples in which monk seal DNA was detected were more abundant in 2019 (n=13, 65% of yearly total) than in 2018 (n=4, 25% of yearly total).

The strongest monk seal DNA recovery within the ferry-route eDNA collection was found in sample MmoMo17 (sample 19-1LiGA1), collected on the 20^th^ of June 2019 at about 11 km from Elba Island and 26 km from Capraia Island, where the monk seal has been sighted at least twice during summer 2020 (June/July). In the same site, exactly one year before (18^th^ of June 2018), a detectable but non-quantifiable signal was identified, witnessing the presence of the monk seal in the Tuscany archipelagos up to two years before the recent *de visu* identifications in the waters surrounding Capraia Island (June-July 2020).

In the Pelagian data, both samples (MmoMc09, and MmoMc11) scoring the strongest detection signals were somehow related to sightings. Sample MmoM+09 was collected in the northern shore of Lampedusa close to where two consecutive sightings would have occurred two months later (Fig. 3). Sample MmoMc11 was instead purposely collected two days after a sighting occurred nearby. In this case, at least for MarVer2, the signal was the strongest within the Pelagian data (5.36*10^3^ DNA copies/L of marine water), comparable in value to the signal obtained in the control eDNA of Madeira sample MmoM+01 (6.38*10^3^ DNA copies/L). We generally found agreement with data reporting a signal persistence of about 48 hours from release (Collins *et al*., 2018), although other factors should be kept in consideration, such as the possible unsubstantiated repeated presence of the animal in the two days between sighting and sampling (which could have increased the signal intensity). Also, we are unaware if the animal/s has/have engaged in behaviors that facilitate the release of biological material (e.g. landing or rubbing on the shoreline), and finally it should be bear in mind the differential eDNA persistence between coastal and deep pelagic waters (Collins *et al*., 2018).

### Relevance of nocturnal samples

This study provides the first example not only of molecular detection of the monk seal presence based on marine water analysis, but it also represents the first case where monk seals are looked for in open waters and, moreover, at any time of day. Despite diurnal (n=17, 47.2%) and nocturnal (n=19, 52.8%) ferry samples were roughly equally represented, positive monk seal DNA detections were more commonly found at night (64.7%, n=11) than in samples collected during light hours (35.3%, n=6). Although we have denominated the ferry samples as “offshore eDNA” (Category 5), the surveyed Livorno-Golfo Aranci route crosses inevitably twice the continental shelf. Thus, 22 (61.1%) of our 36 ferry-based samples were collected off the continental shelf (depth>200 m). These included all the seven samples collected in fix stations 3 and 4 (n=14), plus all the sighting samples (n=8), all occurring along the study route between fix stations 3 and 4 (Fig. 3). About half (n=12, 54.5%) of these 22 deep water samples were collected at night. Interestingly, despite the sample size is not appropriate to establish statistical significance, the instances (n=9) in which monk seal traces were found in offshore waters originated prevalently from nocturnal samples (n=6). These DNA recoveries were however all weak (DBNQ), denoting either diluted (relatively far from the point of shedding) or faded (not freshly released in the environment) signals. Interestingly however, in most of the 22 truly pelagic samples (off the continental shelf) positivity was given by locus MarVer3 (the largest in amplicon size), thus suggesting the presence of not too degraded (probably recently shed) eDNA. If confirmed by future offshore screening, the finding would indicate that monk seals are likely to frequent deep waters -probably foraging-at night, while they have more coastal habits during day time. Interestingly, the metabarcoding analysis on the 2018 ferry-sample subset showed significantly higher vertebrate read counts (>95% attributable to bony fish) in nocturnal-vs-diurnal samples (Valsecchi *et al*., in prep.). The sample recording the highest monk seal signal (MmoMo17) was diurnal, but collected on the continental shelf, at about ten kilometers from two islands. This might explain why this species is more commonly seen by the human eye (thus during day-light hours) in coastal water rather than in offshore waters, which are probably more frequently attended by monk seals at night.

### Value of offshore molecular surveillance over monk seal occurrence

Although recognized as the most endangered pinniped species worldwide, the status of the Mediterranean monk seal population is currently considered “data-deficient” as, so far, its study has been focused on resting/reproductive coastal areas, where paradoxically the research is necessarily limited by the concern of disturbing such a vulnerable species, thus the adoption of camera traps. Yet, beside the above-mentioned constraints, limiting the study of the monk seal to its coastal occurrence would inevitably limit and bias our knowledge of the species, as it brings insights on limited aspects of its life cycle. The possibility to monitor the species in offshore waters opens the prospect to fill the knowledge gaps about still uninvestigated facets of the biology of this threatened species, such as feeding habits, movements during non-reproductive season, species boundaries etc.

### Relevance of the molecular detection approach to the monk seal conservation

There are at least seven circumstances in which molecular monitoring by means of species-specific molecular assays, such as those described here, can find a valid application (Fig. 4):

**Fig.4.**
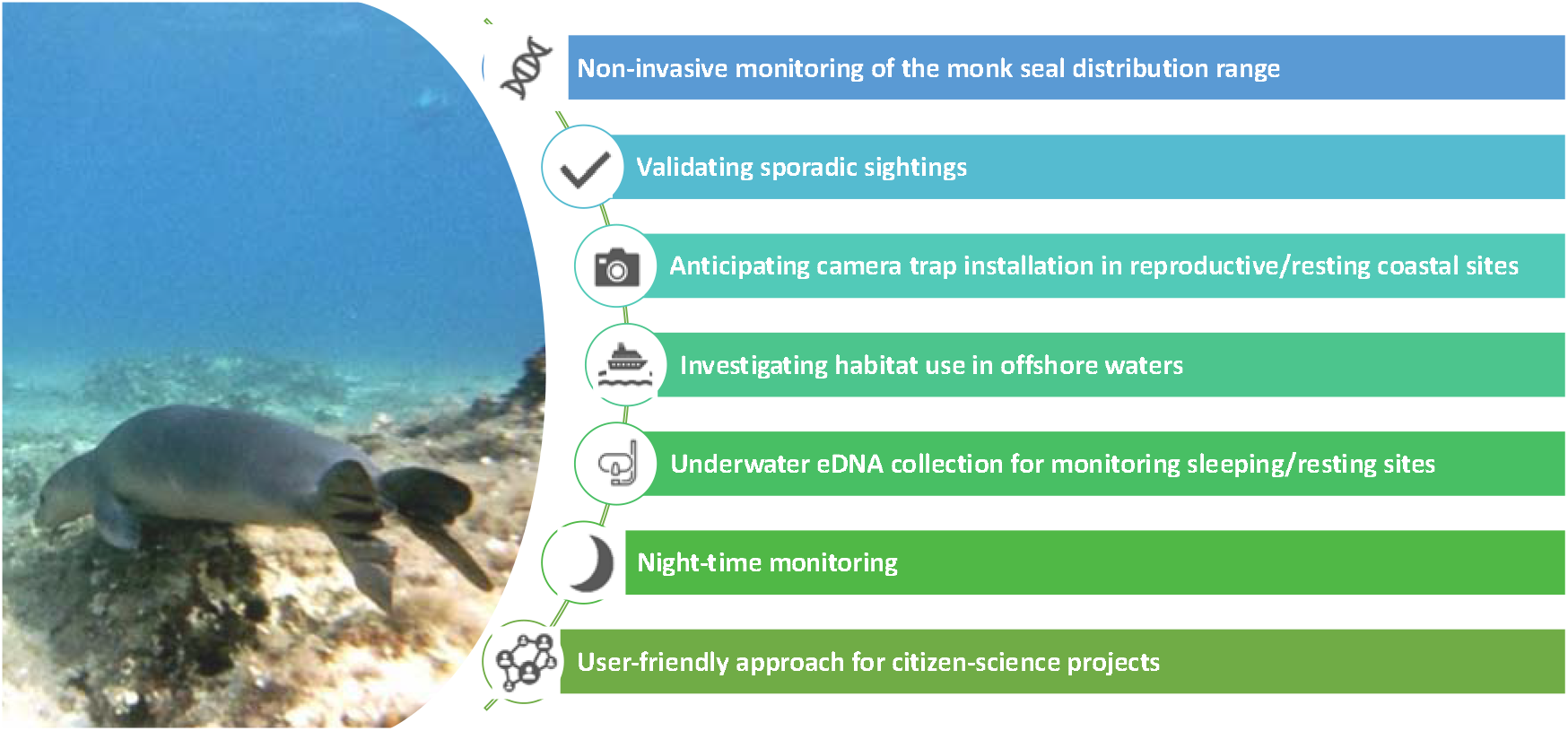
Monk-seal molecular monitoring applications. Possible applications of monk-seal molecular monitoring (photo by Emanuele Coppola)

1. non-invasive monitoring of areas where the presence of the monk seal is already known, in order to estimate their seasonal use and the degree of site fidelity. Visual observations are particularly difficult during the winter seasons, unlike what can be “observed” through molecular monitoring.
2. validating sporadic sightings, on the occasion of which it would be possible to intensify the sampling in waters adjacent to the encounter, in order to verify whether the sighting is accidental or indicative of an expansion of the distributional range.
3. in the coasts adjacent to the known distribution range periodically monitoring the areas which, due to their physical conformation, constitute the potential habitats of choice for the monk seal, such as cave sheltered from sea disturbance and with possibly an inner beach ideal for resting and/or breeding. Besides helping assess the actual home/distribution range, in these cases, the proposed technique could be used to prelude camera trapping, by identifying the best cave/beach/spot where positioning camera traps. Camera traps have the advantage of being able to observe seals’ behavior (and number), but would be extremely expensive to place where the presence of the seal has not been already documented by either visual or molecular observations.
4. exploring vast open expanses of sea in order to investigate still completely unknown aspects of the life of these charismatic species on the high seas. Thus, the proposed assays could be routinely included in future offshore Mediterranean marine eDNA campaigns.
5. water-sample collection can be carried out also underwater, allowing the monitoring of sites used by monk seals for sleep in apnea.
6. the possibility of night monitoring in all the above circumstances.
7. citizen science is becoming an important mean for the collection of easily accessible environmental data, meanwhile increasing the wide public sensibility to conservation issues. Such an approach was not suitable, so far, to monk seal research, as the predominant conservation attitude is minimizing disturbance to this threatened species. The proposed tool, operable in “remote mode”, far from the seal presence could be easily extended to the community of boaters and recreational sea-goers at least for sample collection.

In conclusion, we present three molecular assays tailored for the detection of monk seal DNA, validated on three sets of positive control samples, including eDNA samples collected during the 2020 monk seal breeding season off Madeira (Desertas Islands). The eDNA-based assays have proven to be able to successfully detect the presence of this threatened species in both tested sets of coastal and offshore Mediterranean eDNA, witnessing its presence long before being recorded by human eyes. These outcomes would also suggest that the Mediterranean monk seal species is progressively re-appropriating its original range. Not only could this approach represent an innovative and absolutely non-invasive way to monitor the status of this unique pinniped, but it also has the potential to highlight still uninvestigated aspects of its behavior, such as its nocturnal hunting nature, which is not surprising, but witnessed for the first time thanks to eDNA analysis. More capillary surveys on more extended Mediterranean districts are likely to unveil many other aspects of the biology of this charismatic species. Overall, the presented data encourage the application of an eDNA-based approach as a molecular surveillance tool to be coupled with traditional ecological surveys in order to gather more accurate predictions of species distributions, leading to potentially more effective marine conservation planning.

## ACKNOWLEDGEMENTS

We thank Giuseppe Sciancalepore of the Dept. of Comparative Biomedicine and Food Science of the University of Padova for providing the monk seal archival sample from the Mediterranean marine mammal tissue bank (www.marinemammals.eu). We are grateful to Corsica and Sardinia Ferries, in the person of Cristina Pizzutti, for welcoming our proposal of attempting eDNA sampling from their fleet, thanks to the brokerage of Dr Antonella Arcangeli of ISPRA who set up the FLT network within which our protocol was firstly proposed. Thanks to rangers from the IFCN for the samples collection off Desertas Islands, to Moby Diving and Triton Research for giving support in sample collection off Lampedusa, and Franco D’Urso of the KRCA for promptly supplying BiBSS to Madeira’s team. Thanks to Gruppo Foca Monaca APS - Rome- and Grupa Sredozemna Medvjedica - Zagabira-for providing the biological residues samples from their archives and the data on occasional observations of the last decades. Thank also to University of Bicocca Marine Science past and current master students Roberto Lombardi, Sofia Frappi, Simone Innocente and Francesca Gagliardi for lending aliquots of the samples collected for their theses used to test the assays proposed in this study. Finally, we would like to publicly thank all the supporters of our “MeD for Med” (“Marine eDNA for the Mediterranean”) crowdfunding project, who enthusiastically believed in our initiative despite the fundraising campaign was launched during COVID pandemic, restricting our sampling design.

## DECLARATIONS

## Availability of data and material

Should the manuscript be accepted, the data supporting the results will be archived in an appropriate repository (Dryad, Figshare or Hal) and the data DOI will be included at the end of the article

## Authors’ contributions

Conceptualization: Elena Valsecchi; Methodology: Elena Valsecchi, Antonia Bruno; Formal analysis and investigation: Elena Valsecchi, Antonia Bruno; Writing - original draft preparation: Elena Valsecchi; Writing - review and editing: Emanuele Coppola, Rosa Pires, Andrea Parmegiani, Maurizio Casiraghi; Paolo Galli, Antonia Bruno; Funding acquisition: Elena Valsecchi, Paolo Galli, Maurizio Casiraghi; Resources: Emanuele Coppola, Rosa Pires, Andrea Parmegiani; Supervision: Elena Valsecchi.

## Summary of Supporting Information

**Figure S1 Map of monk seal sightings**

**Table S1 Schematization of sample characteristics and results**

**Figure S2 Log**_**2**_**DNA copies graphs considering all 73 samples**

**Figure S3 Signal drop tissue vs eDNA statistics**

**Figure S4 Hypothesis of “Environmental Marker Selection”**

**Figure S5 Multiplex-test gel**

**Figure S6 Map of MmoM+01 and MmoM+02 sampling sites**

